# *In vivo* multiomic Perturb-seq with enhanced nuclear gRNA capture

**DOI:** 10.64898/2026.03.15.711739

**Authors:** Xinhe Zheng, Jiwen Li, Kwanho Kim, Sean K. Simmons, Ziyan Zhao, Melodi Tastemel, Nhan Huynh, Huixian Qiu, Juntong Ye, Cassandra M. White, Joshua Z. Levin, Xin Jin

## Abstract

*In vivo* CRISPR screening with joint transcriptomic and chromatin readouts has been limited by inefficient recovery of gRNAs from nuclei. Here, we developed *in vivo* multiomic Perturb-seq, an effective platform combining nuclear transcript anchoring with gRNA-specific capture and amplification to enable high-fidelity, high-recovery gRNA assignment and scalable perturbation-resolved single-nucleus multiomics. Applying this platform to interrogate neurodevelopmental disorder risk genes in the developing cortex reveals cell-type-specific transcriptomic and epigenomic perturbation phenotypes.

## Main text

Pooled single-cell CRISPR screens such as Perturb-seq have enabled scalable dissection of gene function by linking genetic perturbations to single-cell transcriptomic phenotypes in heterogeneous cellular populations^1, 2^. Recently, these screens have been extended to capture epigenomic features, such as chromatin accessibility. These multimodal readouts connect transcriptional phenotypes to their underlying chromatin-level regulatory mechanisms, as has been shown by several *in vitro* platforms^3–6^. However, although Perturb-seq has been adapted for *in vivo* use across diverse tissues^7–10^, integrating chromatin accessibility profiling into these workflows has remained challenging. Nucleus-based preparations — required for both *in vivo* tissue dissociation and ATAC-seq chemistry — deplete cytoplasmic gRNA transcripts, reducing recovery below the threshold required for reliable perturbation identity assignment.

To address this gap, we modified adeno-associated virus (AAV)-compatible *in vivo* Perturb-seq^9^ to incorporate three key features. First, to minimize loss of guide RNA (gRNA)-containing transcripts during nuclei isolation, we engineered an RNA hairpin–binding protein system to retain gRNA-derived transcripts at the outer nuclear membrane (**Figure. 1a**)^11^. Second, we designed a gRNA cassette that, analogous to the CROP-seq design^12^, contains U6-driven gRNAs for genome editing and a polyadenylated gRNA-derived transcript that generates a stable and recoverable transcript pool for nuclear anchoring and detection, thereby enabling direct recovery of perturbation identity **(Figure. 1a)**. Third, we adapted the BD Rhapsody multiome workflow for gRNA-specific capture by incorporating custom oligonucleotides complementary to the gRNA-derived transcript on capture beads, together with an optimized nested target-enrichment (TE) PCR step, enabling recovery of gRNA identity together with its linked cell barcode (**Supplementary Figure. 1; Methods)**^13^.

**Figure 1.**
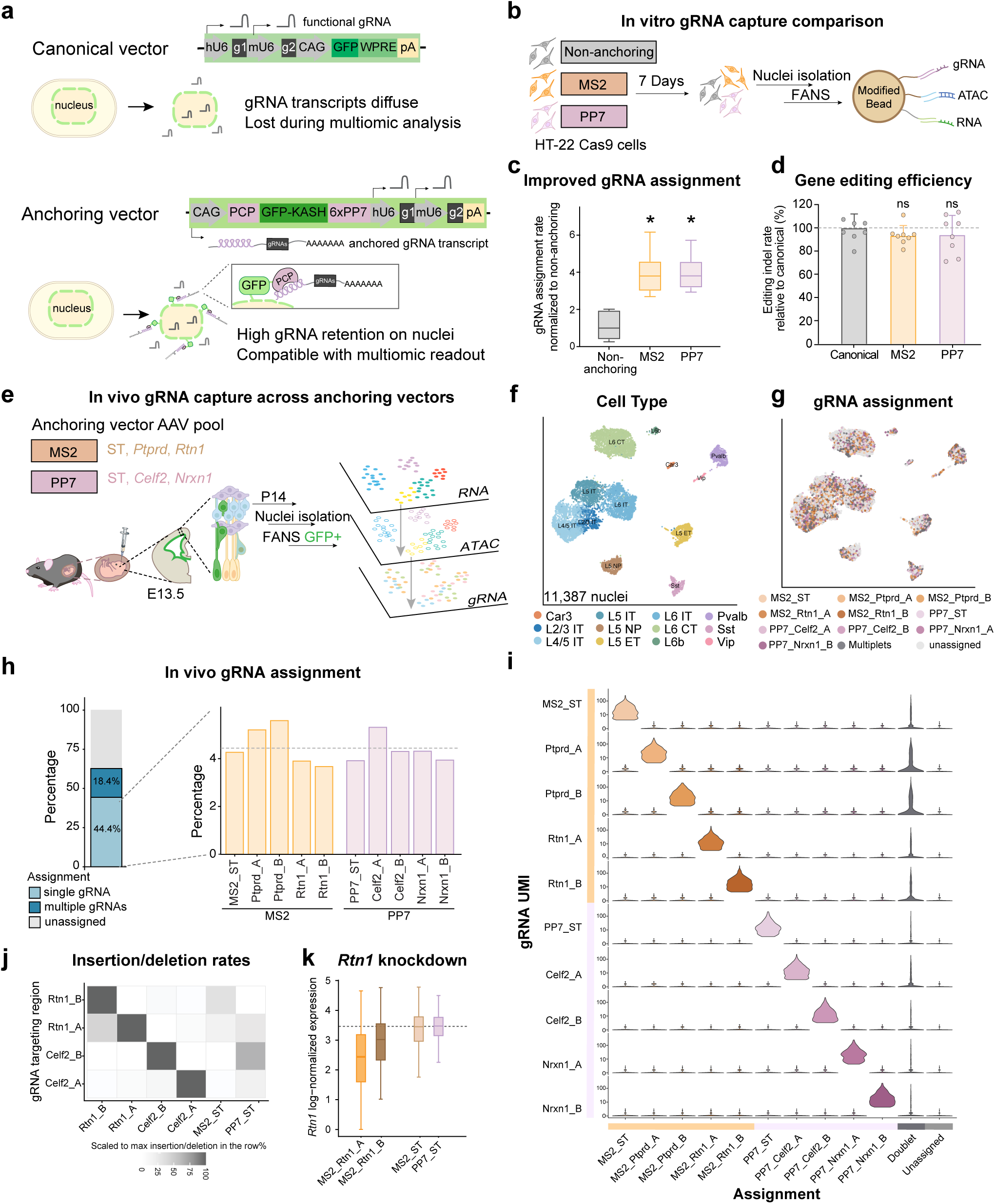
sn-Multiome-compatible nuclear anchoring and targeted capture of gRNA transcripts enable efficient gRNA assignment *in vitro* and *in vivo*. (a) Schematics of vector engineering strategy to improve recovery of gRNA-derived transcripts in nuclei isolation. (b) Experimental workflow for *in vitro* benchmarking of gRNA capture efficiency. (c) gRNA assignment efficiency normalized to the non-anchoring construct. Box plots show the mean (center line), interquartile range (box), and range (whiskers). Asterisks indicate adjusted P values from two-sided Welch’s t-tests (MS2 or PP7 vs. non-anchoring) with Holm correction for multiple testing (*P_adj_* < 0.05). (d) Genome editing efficiency measured *in vitro* across vector designs. A canonical vector (Figure. 1a upper panel) only expresses U6-driven gRNAs for genome editing without gRNA-derived polyadenylated transcripts. Error bars represent standard deviation. Statistical significance was assessed by two-sided Welch’s t-tests (MS2 or PP7 vs. canonical) with Holm correction for multiple testing (ns, not significant). (e) Experimental design for *in vivo* sn-multiome perturbation profiling. (f-g) Uniform manifold approximation and projection (UMAP) visualization based on weighted nearest neighbor (WNN) integration of nuclei profiled *in vivo*, colored by cell type (f) and gRNA assignment (g). ST: safe target. (h) gRNA assignment rates across vectors *in vivo*. The dotted line indicates the average assignment percentage across single gRNAs. (i) Violin plot showing gRNA transcript levels across gRNA assignments. (j) Heatmap showing insertion/deletion frequencies at gRNA target loci from the multiome dataset across gRNA assignments, scaled by the maximum value in each row. (k) Log-normalized expression of *Rtn1* across assignment categories. The dotted line marks the *Rtn1* level in ST as the baseline.

To optimize gRNA assignment efficiency, we first systematically evaluated key features of the perturbation capture workflow *in vitro* in a single-nucleus multiome (sn-multiome) assay. We benchmarked gRNA recovery using AAV pools containing two RNA hairpin-binding protein pairs (MS2 or PP7) gRNA constructs, compared to the non-anchoring gRNA construct at low multiplicity of infection (MOI) into HT-22 Cas9 cells (**Figure. 1b, Supplementary Figure. 2a-b**). Across constructs, we found that increasing the custom-oligo modification from 20% to 80% on the barcoded single-cell beads (from BD Rhapsody) alone increased single-gRNA assignments by 2.3-fold (**Supplementary Figure. 2c-d**); in addition, TE PCR outperformed *in vitro* transcription (IVT) by an additional 2.7-fold in single-gRNA assignment (**Supplementary Figure. 2d**). In the optimal combination, both MS2- and PP7-anchoring vectors consistently improved gRNA assignment by 3.8-fold relative to non-anchoring vectors (**Figure. 1c**). Importantly, these engineered vectors preserved editing efficiency comparable to canonical vectors across multiple targets (**Figure. 1d**). Together, these results indicate that nuclear anchoring markedly improved recovery of perturbation identity without compromising genome editing efficiency.

**Figure 2.**
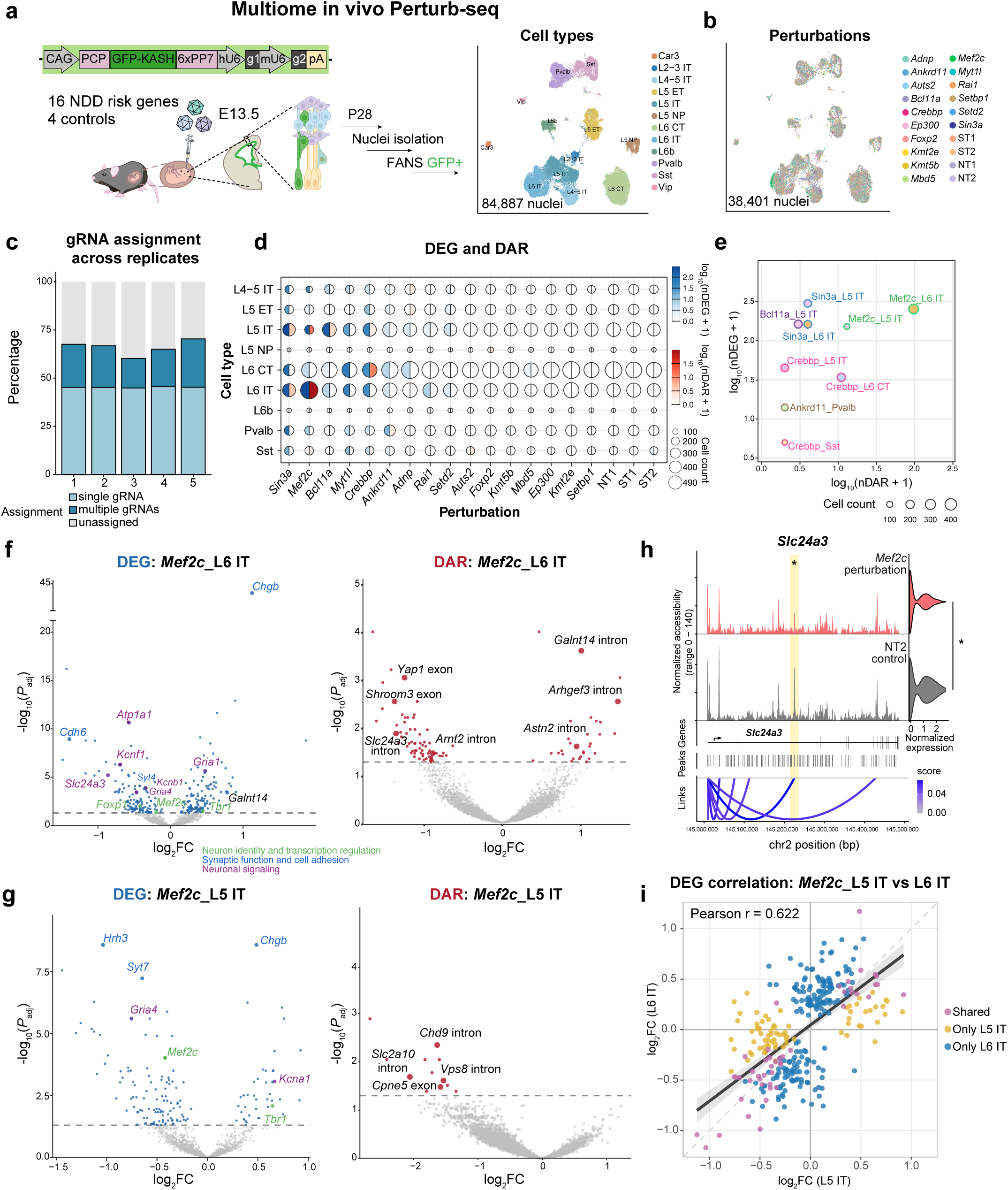
***In vivo* multiome Perturb-seq identified cell-type-specific perturbation phenotypes.** (a) Experimental design for 16-gene *in vivo* multiomic Perturb-seq, and WNN UMAP visualization of all high-quality nuclei, colored by cell type. Nine glutamatergic excitatory populations (L2-3 intratelencephalic (IT), L4-5 IT, L5 IT, L6 IT, Car3, L5 extratelencephalic (ET), L5 near-projecting (NP), L6 corticothalamic (CT) and L6b) and three GABAergic inhibitory populations (Pvalb, Sst, and Vip) were identified. (b) WNN UMAP visualization of all high-confidence single gRNA assigned nuclei colored by perturbation identity. (c) Bar plot showing gRNA assignment rates across 5 replicates. (d) Dot plot of the number of differentially expressed genes (DEGs) and differential accessibility regions (DARs) across cell types and perturbations (*P_adj_* <0.05). For each dot, the color on the left is based on the number of DEGs, the color on the right is based on the number of DARs, and the size is based on the number of nuclei. (e) Scatter plot of DEG and DAR changes in highly impacted cell types and perturbations. Filling colors indicate cell types, and the stroke colors indicate perturbation. Only cell type and perturbation combinations with at least 1 DAR and 1 DEG are shown. (f-g) Volcano plots of cell-type-specific effects for DEGs and DARs by *Mef2c* perturbations in L6 IT (f) and L5 IT (g). Larger dots label example DEGs. (h) Comparison between *Mef2c* perturbed cells and NT2 control cells for gene expression level and chromatin accessibility at *Slc24a3* in L6 IT neurons. Asterisks mark significant DAR, also highlighted in yellow, and DEG (*P_adj_*<0.05). At the top are coverage plots of ATAC-seq fragments along the length of the gene for each perturbation, on the right is a violin plot showing the expression (CP10K) of *Slc2413* in each perturbation. Also shown are the gene structure, the locations of peaks, and a graph linking peaks to genes that have significant correlation. (i) Scatterplot comparing DEG results between L5 IT and L6 IT for *Mef2c* perturbation. The dark-gray line and light-gray shaded band show the linear regression fit and 95% confidence interval computed from all plotted genes, respectively.

To evaluate gRNA assignment *in vivo*, we delivered MS2- and PP7-anchoring AAV vectors into the lateral ventricles of embryos at E13.5, and isolated GFP^+^ cortical neuronal nuclei at postnatal day 14 (P14) for joint single-nucleus RNA, ATAC, and gRNA profiling (**Figure. 1e, Supplementary Figure. 3a**). Across 11,387 high-quality nuclei, 62.8% were assigned with perturbations and 44.4% received a single-gRNA assignment; MS2 and PP7 performed similarly across cell types (**Figure. 1f-h, Supplementary Figure. 3b, 4a**). Our gRNA assignment showed low cross-gRNA correlation, with single-gRNA signals clearly resolved across nuclei, reflecting high signal specificity under the low MOI AAV delivery. (**Figure. 1i, Supplementary Figure. 4b-c**). Overall, gRNA assignment rates from the sn-multiome were comparable to those of single-cell and single-nucleus RNA-seq, representing a marked improvement in sn-multiome gRNA recovery (**Supplementary Figure. 4d**)^9, 14, 15^. Similar to the *in vitro* experiment, 80% bead modification with TE amplification yielded the most robust assignment performance *in vivo* (**Supplementary Figure. 4e**), while RNA and ATAC quality metrics remained comparable across bead modification conditions at similar sequencing depth (**Supplementary Figure. 3c-d, Supplementary Table 2**).

As an orthogonal validation of the gRNA assignment, we quantified insertion/deletion rates at gRNA-targeted loci directly from the transcriptome in sn-multiome data. Assigned nuclei were enriched for target-matched editing signatures, with little cross-gRNA signal (**Figure. 1j, Supplementary Figure. 4f**). Nuclei carrying *Rtn1*-targeting gRNAs showed reduced *Rtn1* expression as expected, supporting the accuracy of our assignments (**Figure. 1k**).

We next applied this sn-multiome Perturb-seq platform to interrogate perturbation effects of disease risk genes in the developing cortex. We designed a pooled screen targeting 16 transcription factors and chromatin regulators implicated in neurodevelopmental disorders (NDDs) that are broadly expressed during corticogenesis across multiple neuronal cell types^16–18^ (**Supplementary Figure. 5**)^19^. The equally pooled AAV library was delivered into the lateral ventricles of Cas9-expressing embryos at E13.5 with low MOI, and GFP^+^ cortical neuronal nuclei were profiled at P28 using our sn-multiome workflow (**Figure. 2a, Supplementary Figure. 6a-c**). Across 15 animals from 5 experiments, we recovered 84,887 high-quality nuclei spanning 12 neuronal cell types, including nine glutamatergic excitatory populations and three GABAergic inhibitory populations (**Figure. 2a; Supplementary Figure. 7a-g)**. Overall, 66.2% of these cells were assigned to perturbations and 45.2% with single-gRNA assignments, yielding 38,401 nuclei for downstream analysis, consistent with the scalability of AAV-based Perturb-seq outperforming lentivirus *in vivo*^9^ (**Figure. 2b-c, Supplementary Figure. 7b**). Assignment performance was consistent across 5 biological replicates (**Figure. 2c**), and each perturbation was represented by 1,516-2,361 nuclei (**Supplementary Figure. 7c**).

To define perturbation-induced molecular phenotypes, we performed pseudobulk differential expression gene (DEG) and differential accessibility region (DAR) analyses for each perturbation-by-cell-type pair. Comparisons among control pairs yielded minimal DEG and DAR signals (<=1 DEG and no DARs), confirming low false discovery rates (**Figure. 2d**). Across the screen, the number of DEGs and DARs identified for each perturbation differed markedly across cell types; phenotypic changes were detected in gene expression, chromatin accessibility, or both (**Figure. 2d-e**). Perturbations of *Myt1l*, *Rai1*, and *Setd2* produced phenotypes detected primarily through transcriptional changes, whereas perturbations of *Sin3a*, *Mef2c*, *Bcl11a*, *Crebbp*, and *Ankrd11* produced substantial effects in both modalities **(Figure. 2d-e)**.

Among perturbations with substantial effects in both gene expression and chromatin accessibility, the resulting molecular phenotypes were strongly cell-type-specific. Loss of *Sin3a*, a chromatin-associated transcriptional coregulator, produced broad DEGs and DARs across multiple excitatory and inhibitory neuronal cell types (**Figure. 2d, Supplementary Figure. 8a**). By contrast, transcription factor *Mef2c* showed cell-type-restricted multimodal phenotypes, with prominent changes detected in L5 IT (151 DEGs and 12 DARs) and L6 IT neurons (253 DEGs and 96 DARs); it has minimal effects in deep-layer excitatory neuron subtype L6 CT (4 DEGs and 0 DARs), though with similar sampling power (**Figure. 2d-g**). In L5 IT neurons, *Mef2c* perturbation showed decreases in both gene expression and chromatin accessibility, consistent with a concordant shift in regulatory state (**Figure. 2g**). In L6 IT neurons, joint

RNA-ATAC profiling further linked accessibility changes to expression changes at specific loci: *Slc24a3*, *Rimbp2, Arnt2* and *Galnt14* showed concordant changes in both accessibility and expression level (**Figure. 2h, Supplementary Figure. 9**). At *Slc24a3* locus, accessibility of a DAR was correlated with *Slc24a3* expression, suggesting a putative cis-regulatory interaction. In addition, *Mef2c* perturbation affected genes linked to neuronal identity and transcriptional regulation (*Tbr1, Foxp1*), synaptic function and cell adhesion (*Chgb, Cdh6, Syt4*), and neuronal signaling (*Atp1a1, Kcnb1, Kcnf1, Gria1, Gria4*) (**Figure. 2f**). We compared these phenotypes with a published *Mef2c* conditional knockout bulk RNA-seq dataset and observed correlated fold change of shared DEGs, validating our results (Pearson r = 0.816) (**Supplementary Figure. 8b**)^20^. Although *Sin3a* and *Mef2c* are both broadly expressed across neuronal subtypes (**Supplementary Figure. 5**), their perturbation effects markedly differ, indicating baseline expression alone may not be sufficient to predict their functional impact in heterogeneous cell types *in vivo*. Cell-type-restricted patterns further showed that transcriptional responses to *Sin3a* perturbation (321 DEGs, Pearson r = 0.800) were more similar between L5 IT and L6 IT than responses to *Mef2c* perturbation (312 DEGs, Pearson r = 0.622) (**Figure. 2i, Supplementary Figure. 8c**). The cross-cell-type DEG overlap patterns likewise indicated broader sharing of *Sin3a*-induced signatures but greater divergence of *Mef2c*-induced signatures across cell types, demonstrating that our platform can effectively resolve both broadly shared and subtype-restricted perturbation signatures with high fidelity (**Supplementary Figure. 8d-e**).

By simultaneously linking perturbation-induced transcriptional changes to their chromatin accessibility from the same cell, *in vivo* multiomic Perturb-seq could enable construction of more complete, cell type-resolved models of gene regulatory networks. Measuring gene expression and chromatin accessibility in the same perturbed cell could provide mechanistic resolution that is difficult to achieve with unimodal assays and critical for understanding disease-associated mechanisms *in vivo*^21, 22^. Because our strategy is compatible with AAV-based delivery and nucleus-based multiomic workflows, it is broadly applicable across tissues, species and disease models. More generally, this framework provides a foundation for building predictive models of gene function from multimodal perturbation data in a cell type-specific manner, enabling in silico prediction of more generalizable inference of gene function and regulatory logic *in vivo*.

## Methods

### gRNA design and vector construction

For data related to Figure. 1b, e and 2a: gRNA were designed using CRISPick^23, 24^ and a pair of gRNAs targeting >100 bp apart were chosen manually, prioritizing targeting early coding exons. We also included non-target (NT) controls that do not target anywhere in the mouse genome and safe-target (ST) controls that do not target coding genes in the mouse genome. For data related to Figure. 1j: For a sanity check of gRNA assignment, gRNAs targeting the 3’UTR region were designed using an online tool at <u>benchling.com</u> (CRISPR function), such that the target sites would lie in regions with higher coverage in sn-multiome sequencing, allowing detection of CRISPR-induced indels. All the gRNAs used in this work are listed in ***Supplementary Table 1***. AAV gRNA vectors were cloned in array format by Golden Gate and confirmed by Sanger sequencing^25^. Plasmids generated in this study will be deposited to Addgene (https://www.addgene.org/Xin_Jin/).

### Viral vector production and quality control

AAV production and titration were performed according to the published protocol^25^. For the pooled AAV gRNA library, each gRNA plasmid was transfected into a separate dish of HEK293 cells, and the harvested supernatants and cell pellets were combined prior to AAV purification. To assess preservation of gRNA representation in the AAV pool, a gRNA distribution quality control assay was performed on the final purified AAV pool. Briefly, an aliquot of AAV was used to remove residual unpackaged plasmid DNA, followed by heat inactivation and proteinase K digestion to release encapsidated viral genomes. Viral DNA was used as the template for PCR amplification of the dual-gRNA cassette (primers see ***Supplementary Table 1***). Amplicons were quantified and sequenced on a nanopore platform (Plasmidsaurus premium PCR service) to a depth sufficient to achieve >100 reads per gRNA on average. Reads were aligned to the reference gRNA library with a custom Python script using the expected amplicon structure and guide reference sequences (see ***Code Availability***). Briefly, a read was counted only when both gRNA1 and gRNA2 were confidently detected at the expected positions. Reads containing the designed gRNA1-gRNA2 combinations were classified as on-target pair assignments.

### Cell culture and viral transduction

Mammalian cell culture experiments were performed in the HT22-Cas9 mouse hippocampal neuronal cell line (Ubigene, YC-A004-Cas9-H) grown in DMEM (Thermo Fisher Scientific, #11965092) with 25 mM glucose, additionally supplemented with 1× penicillin–streptomycin (Thermo Fisher Scientific, #15140122), and 5-10% fetal bovine serum (Thermo Fisher Scientific, #16000069). HT22 cells were maintained at confluency below 80%. For AAV transduction, a MOI of 500 - 2000 was used to achieve <5% GFP^+^ transduced cells. For the vector comparison, cells transduced with different vectors were pooled together for even gRNA distribution based on flow cytometry analysis for GFP^+^%.

### In utero AAV vector administration

All animal experiments were performed according to protocols approved by the Institutional Animal Care and Use Committees (IACUC) of The Scripps Research Institute. All mice were kept in standard conditions (a 12-h light/dark cycle with *ad libitum* access to food and water). AAV (0.5-1.5 µL per embryo, with AAV stock concentration at 1×10^12^-9×10^13^ Units/mL) was administered in utero to the lateral ventricle at E13.5 in CD1 or Cas9 transgenic mice (Jax, #026179) for immunohistochemistry analysis and sn-multiome sequencing.

### Immunofluorescent staining

Postnatal pups (P14 and P28) were anesthetized and transcardially perfused with ice-cold PBS followed by ice-cold 4% paraformaldehyde in PBS. Dissected brains were postfixed overnight in 4% paraformaldehyde at 4 °C. Postnatal brains were embedded in 2% agar and 80 μm tissue sections were collected on a vibratome. The slides with mounted tissue sections were washed 4 times with PBS with 0.3% TritonX-100 and incubated with blocking media (10% donkey serum (Sigma Aldrich, #S30-100ML) in 0.3% TritonX-100 with PBS) for 2 hours at room temperature, then incubated with primary antibodies in the blocking media overnight at 4°C. Slides were washed with PBS with 0.3% TritonX-100 4 times. Secondary antibodies were applied at a 1:1000 dilution in blocking media and incubated for 2 hours at room temperature. Slides were then washed 4 times with PBS with 0.3% TritonX-100 and incubated with DAPI (VWR, #76482-848, 1:5000) for 10 mins before mounting with Antifade Mounting Medium (Vector Laboratories, #H-1700-10). All images were taken using a Nikon AX Confocal Microscope with a 10x air or 20x air objective.

The primary antibodies and dilutions were: Rabbit anti-GFP antibody (Invitrogen, #A-11122, 1:500) Goat anti-RFP antibody (Rockland, #200-101-379, 1:500), FluoTag-X2 anti-TagFP AlexaFluor 647 (NanoTag Biotechnologies, #N0502-AF647, 1:200).

### Insertion/Deletion analysis in vitro

HT22-Cas9 cells in a 96-well plate were transfected with 200 ng gRNA expressing plasmid using Polyethylenimine HCl MAX, linear, MW 40,000 (PEI, Polysciences, #24765-1). After 36 hours, genomic DNA was extracted using QuickExtract (New England Biolabs, #B6058S). Target loci spanning the expected Cas9 cut sites were PCR-amplified using a two-step PCR strategy to append Illumina adapters and sample indices. Indexed amplicons were pooled, gel-purified, and quantified by qPCR prior to sequencing on iSeq100 (amplicon-specific primers see ***Supplementary Table 1***). Demultiplexed FASTQ files were used to determine editing outcomes by quantifying the frequency of insertions and deletions within a defined window centered on the cut site relative to the unedited reference amplicon sequence using CRISPResso2 v.2.2.15^26^.

### Gene expression enrichment in developing cortex

Gene expression analysis was performed using the mouse visual cortex development cell type atlas^19^. The dataset was subsetted to include 16 predefined target genes, and cell types that were not targeted in the screen (OPC-Oligo, Vascular, Astro-Epen, and Immune) were excluded based on the class_label annotation. The gene expression dot plot was generated using Scanpy v1.11.2^27^ (sc.pl.dotplot). For each gene within each group, dot size indicates the fraction of cells with expression >0 (expression_cutoff=0.0), and dot color indicates the mean expression across cells in that group; color values were scaled per gene across groups for visualization (standard_scale=“var”). Additional analyses were conducted by subsetting the dataset to E13.5 and P28 timepoints to produce subclass-by-gene dotplots.

### Single-nucleus multiome

*<u>Nuclei isolation:</u>* P28 mouse cortices were rapidly dissected and flash-frozen in liquid nitrogen prior to nuclei isolation. Nuclei isolation medium (250 mM sucrose, 25 mM KCl, 10 mM HEPES, pH 7.5, and 5 mM MgClL in nuclease-free water) was prepared up to 24 h in advance and stored at 4 °C. Lysis buffer (nuclei isolation medium supplemented with 1 mM DTT, 0.05% NP-40, 1% BSA, 10 mg/mL Kollidon VA64 (Sigma-Aldrich, #190845), and 0.6 U/µL RNase inhibitor (Fisher Scientific, #NC1081844)) and washing buffer (nuclei isolation medium supplemented with 1 mM DTT, 1% BSA, and 0.6 U/µL RNase inhibitor (Fisher Scientific, #NC1081844)) were prepared fresh on the day of use. Frozen tissue was homogenized on ice in 1 mL lysis buffer using a pre-chilled dounce homogenizer. Homogenization conditions were optimized to achieve complete dissociation without visible tissue fragments. An additional 1.5 mL lysis buffer was added, and the homogenate was incubated on ice for 3 min. The lysate was filtered through a 70-µm cell strainer, diluted with washing buffer, and centrifuged at 200 × g for 5 min at 4 °C. The nuclei pellet was resuspended in washing buffer and incubated with anti-NeuN–AF647 antibody (Abcam, ab190565, 1:500) and mouse nucleoporin P62 sample tag antibody (BD, 460293-460297, 1:200, only for 16-gene screen) for 15 min on ice.

For nuclei isolation of HT22 cells, cells were washed twice with cold DPBS, scraped with 5mL lysis buffer (nuclei isolation medium supplemented with 1 mM DTT, 0.2% Tween-20, 0.2% IGEPAL CA-630, 1% BSA and 0.6 U/µL RNase inhibitor (Fisher Scientific, #NC1081844)), and incubated on ice for 5 min, diluted with washing buffer, and centrifuged at 300 × g for 5 min at 4°C.

After two washes (150 × g, 8 min, 4 °C), nuclei were resuspended in the washing buffer and stained with 7-Aminoactinomycin D (7-AAD, Thermo Fisher Scientific, #A1310). Perturbed nuclei were enriched by fluorescence-activated nuclei sorting (FANS) to enrich 7-AAD^+^/GFP^+^/NeuN^+^ neuronal nuclei using MoFlo Astrios Cell Sorters (Beckman Coulter) or a Bigfoot™ Spectral Cell Sorter (Thermo Fisher Scientific) with 100 µm nozzle.

*<u>Single-nucleus multiome library construction and quality control:</u>* The sn-multiome libraries were constructed following BD Rhapsody Single-Cell ATAC-Seq BD OMICS-One™ WTA Next and Sample Tag Library Preparation Protocol (23-25007(01)) with modifications to construct the gRNA library. BD Rhapsody beads were modified as described in^13^ to add a splint oligo to capture tagmented genome. Sorted nuclei were immediately concentrated using activated 1 x CUTANA Concanavalin A Conjugated Paramagnetic Beads (EpiCypher, #21-1401) as described in^28^ and resuspended in modified nuclei buffer. 30-60k nuclei were tagmented and loaded to a BD Rhapsody cartridge; modified BD beads were loaded to the cartridge, lysed, and Rhapsody beads with cDNA and tagmented DNA were retrieved from the cartridge. Rhapsody beads were processed with lysis, ligation, reverse transcription, splint oligo removal, and Exonuclease I treatment, and stored at 4 °C. The ATAC library was constructed following the BD Rhapsody protocol. Exonuclease I-treated beads were subjected to gRNA library construction with modification from BD Rhapsody targeted enrichment mRNA protocol (23-24121(03)). Exonuclease I-treated beads were then processed with the mRNA library and sample multiplexing tag library construction.

### Sequencing

The gene expression library, gRNA library and sample multiplexing tag library were sequenced with a NextSeq 2000 100-cycle kits or a NovaSeq X 100-cycle kit (Illumina) with sequencing saturation to ensure greater than 20,000 reads (RNA), 1,000 reads (gRNA) and 1,000 reads (sample tag) coverage per nucleus, respectively (R1: 63 cycles, R2: 67 cycles, i7: 8 cycles). The ATAC library was sequenced with a NextSeq 500 or a NovaSeq X 200-cycle kit (Illumina) with sequencing saturation to ensure greater than 25,000 reads per nucleus (R1: 60 cycles, R2: 60 cycles, i7: 8 cycles, i5: 60 cycles).

### Single-nucleus multiome data preprocessing and QC filtering

BCL files were used to generate FASTQ files using the default parameters by “bclconvert” command (v3.10.5). The BD Rhapsody Sequence Analysis Pipeline v3.0 was used for alignment of RNA and ATAC libraries to the mm39 reference genome (prebuilt reference package from BD version 2023-09). Demultiplexed reads for RNA (including mRNA, gRNA and sample tag (for the 16-gene screen)) and ATAC libraries were input into the pipeline using a YAML file with default parameters. Expected cell number and supplemental reference of gRNA targeted sequences in a FASTA format were added to the YAML file. Sample tag version was additionally indicated for the 16-gene *in vivo* dataset. Both RNA and ATAC data were used to predict cells (Cell_Calling_Data = mRNA_and_ATAC), using the default algorithms. Expected cell counts were predicted for each run, taking 30% from the number of nuclei sorted. The Generate_Bam parameter was set to “true” for cell level quality control.

For RNA data processing, unfiltered counts matrices were loaded into R (v4.4.3) using the Read10X function from the Seurat R package (v5.3.0)^29^. Seurat object assay version was set to v3 (options(Seurat.object.assay.version = “v3”)). The Seurat object was created from each of the count matrices (CreateSeuratObject with min.cells=10 and min.features=200) and subsequently combined into a single Seurat object. Transgene UMI counts were removed from the gene counts matrix and stored separately in the “gRNA” assay of the Seurat object. For cell-level QC, in addition to standard metrics, CellLevel_QC (https://github.com/seanken/CellLevel_QC) was run using the individual BAM files. Cells were also scored to identify potential doublets within their own batches using the scds R package (v1.13.1 in R v4.2.2)^30^. The data were then normalized (NormalizeData), scaled on variable genes (FindVariableFeatures) while regressing out total UMI counts and percent mitochondrial gene expression (ScaleData with vars.to.regress = c(“nCount_RNA”, ”percent.mito”)), and reduced in feature dimensions (RunPCA). The top 30 PCs were used to further reduce dimensions for visualization (RunUMAP with dims=1:30), and to identify neighboring cells (FindNeighbors with dims=1:30). Cell clusters were identified using FindClusters function with resolution=0.5. Combining information from QC metrics, low-quality clusters (characterized by relatively low numbers of both UMIs and genes (median gene count < 1,000), high percent mitochondrial gene expression (median percent mitochondrial gene expression > 8), or a higher rate of reads aligned to intronic regions (median fraction of reads in intronic region > 0.55)) were identified and removed. Low-quality cells were also excluded from the analysis if they possessed fewer than 500 genes, fewer than 1000 or greater than 15,000 total UMIs, or a doublet score (hybrid_score from scds) > 1.5. For the dataset with sample tags, only the cells with sample tags confidently assigned by BD Rhapsody pipeline were selected. Following QC and filtering, the remaining cells were processed through the same pipeline as described above. To correct for sequencing batch effects in the 16-gene screen, we added an integration step using Harmony (v1.2.4)^31^ after the PCA step (RunHarmony function with group.by.vars set to cells sequenced together) and used the top 20 harmony dimensions for the downstream processes of generating UMAP and identifying cell clusters.

For ATAC data processing, peak matrices, generated by the BD Rhapsody sequence analysis pipeline, were loaded using the Signac R package (v1.15.0)^32^. A common peak set was identified across all samples using the UnifyPeaks function (mode=”reduce”). Peak counts for each cell were generated (FeatureMatrix) and assembled into a ChromatinAssay mapping to the mm39 reference genome. Gene annotations for GRCm39 from EnsDb (id=“ AH119358”) were added using AnnotationHub (v3.14.0). Quality control metrics were calculated for each cell and used to select high quality cells: library size (nCount_peaks) between 200 and 10,000 with at least 10% of reads within peaks and a minimum transcription start site (TSS) enrichment score of 1. Cells with a high proportion of reads in genomic blacklist regions (> 7.5%) or abnormal chromatin packaging (indicated by nucleosome signal > 0.3) were excluded. For the dataset with sample tags, only BD predicted cells with sample tags confidently assigned were used.

### Multiome data processing

Cells identified in both the fully processed RNA Seurat object and the QC-filtered ATAC object were retained to establish a common cell set. The ATAC peak matrix and its corresponding quality control metadata were then added to the RNA Seurat object as a separate assay. For the ATAC data, the peak counts were normalized using term frequency-inverse document frequency (TF-IDF) via the RunTFIDF function of the Signac R package. Variable features (top peaks) were selected using the FindTopFeatures function with a minimum cutoff set to ’q0’. Dimensionality reduction was then performed using singular value decomposition (SVD) via the RunSVD function. To visualize the ATAC data independently, RunUMAP function of Seurat was run on dimensions 2 through 30 of the latent semantic indexing (LSI) reduction, explicitly excluding the first dimension as it typically correlates with sequencing depth. To integrate both modalities into a unified representation, a Weighted Nearest Neighbor (WNN) graph was constructed using the FindMultiModalNeighbors function from Seurat (dims.list=list(1:50, 2:50)). A joint WNN UMAP was generated based on this weighted nearest neighbor graph (RunUMAP with nn.name=”weighted.nn”). Final multimodal cell clustering was performed directly on the WNN graph (graph.name=”wsnn”) with a clustering resolution of 0.3 (FindClusters).

### gRNA assignment

Cellular assignment to gRNA perturbations was performed using the Crispat Python package (v0.9.10 in Python v3.10.19). Raw gRNA count matrices were converted to AnnData objects, and cells were assigned to their respective gRNAs using a Gaussian mixture model approach (crispat.ga_gauss with inference=’vi’) with a minimum UMI threshold of 3. Cells with no gRNA assigned were labeled “Not assigned” and those with multiple gRNAs were labeled “Doublet”.

### Cell type annotation

To annotate cell types, the MapMyCells web portal (RRID:SCR_024672)^33^ from the Allen Brain Institute was used. The RNA Seurat object was first downsampled to a maximum of 500 cells per cluster. Raw RNA counts were extracted, mouse gene symbols were converted to ENSEMBL IDs, and duplicated genes were removed before exporting the dataset into an AnnData format (.h5ad) for upload. The data were queried against the 10x Genomics Whole Mouse Brain reference (CCN20230722) using the Hierarchical Mapping algorithm. The resulting mapping classifications—including class, subclass, and cluster names, as well as cluster bootstrapping probabilities—were integrated back into the Seurat object cell metadata. This prediction, combined with cluster-specific DEGs, was used to annotate clusters with cell type labels.

### Differential expressed gene (DEG) and differential chromatin accessibility region (DAR) analysis

For DEG analysis, each cell type was analysed one by one. A pseudobulk RNA count matrix was created, combining all cells with the same guide assignment and sample tag assignment, using the AggregateExpression command in Seurat v4.0.0 with the slot=”counts” argument. Genes that did not have >10 UMI in at least 2 samples were excluded. Using EdgeR v3.32.1^34^, a DGEList object was created and normalization factors were calculated, then filterByExpr was used to filter out lowly expressed genes. The dispersions were estimated with estimateDisp, and a model was fit with glmFit, using a design matrix that had both guide RNA assignment and sample tag assignment as covariates. The glmLRT function was used to perform the actual statistical test on each perturbation (comparing each perturbation to the NT2-g2 control), and Benjamini Hochberg was performed with the p.adjust command.

For differential chromatin accessibility analysis the same approach was used as for the differential expression analysis, with two changes. First, instead of pseudobulking the RNA count matrix, the peak count matrix was used to create a pseudobulk. Second, for filtering the filterByExpr function was not used.

For linking DEG and DAR results, the LinkPeaks function in Signac was used to find correlation based links between genes and peaks, and the CovaregePlot function was used to plot the results, with a window of size 2000 bp.

### Insertion and Deletion (indel) ratio analysis from single-cell dataset

To validate gRNA assignments and assess genomic editing efficiency, indels were analyzed directly from sequencing BAM files using a custom Python script relying on the pysam package (v0.16.0.1 in Python v3.7.12) ^35, 36^. Reads were analyzed to determine if they contained an insertion or deletion fully spanning the predefined target regions for each gRNA (fully_spanning=True). Indel ratios for each assigned cell cluster were calculated as the total number of insertions and deletions divided by the total number of fully spanning reads mapping to the given target region.

## Data Availability

The data generated in this study will be submitted to NCBI Gene Expression Omnibus, the Broad Institute single cell portal, and the UCSC cell browser through the NIH SSPsyGene consortium.

## Code Availability

The analysis pipeline will be deposited in the GitHub repository (https://github.com/jinlabneurogenomics/multiome-perturbseq).

## Acknowledgements

We thank David Shin and Tom Nowakowski for their advice; Jixiang Zhang for assistance with method development experiments; Scripps DAR and Flow Cytometry Core for assistance; and all members of the Jin lab for their support. X.Z. was supported by Dorris Scholar Award and the Frank J. Dixon Fellowship. J.L. was supported by CIRM Training Fellowship (EDU4-12811) and Dorris Scholar Award. C.M.W. was supported by the Skaggs-Oxford Program and The Schimmel Family Endowed Fellowship. J.Z.L. was supported by the Stanley Center for Psychiatric Research at the Broad Institute of MIT and Harvard. J.Z.L and X.J. were supported by the National Institute of Health (R01MH137042) and are part of the SSPsyGene Consortium. X.J. and this work were supported by HHMI, Simons Foundation Autism Research Initiative Sex Differences Collaboration (SFARI #736613), CIRM ReMIND-L DISC4-16295, National Institute of Health (R01HG012819, R01AT013748), the Mathers Foundation, the One Mind Foundation, the Conrad Prebys Foundation, the Pew Charitable Trust, the McKnight Endowment, the Astera Institute, Jed McCaleb, and James Fickle.

## Author Contribution

X.Z. and X.J. conceived the study with input from all authors. X.Z., Z.Z. and C.M.W. performed molecular cloning. N.H. and H.Q. prepared AAV vectors. X.Z. performed sn-multiome experiments with help from J.L., M.T., and C.M.W. X.Z., S.K.S., and K.K. analyzed the Perturb-seq data under the supervision of J.Z.L. and X.J. J.Y. analyzed gene-expression data from published datasets. X.Z., J.L., and X.J. drafted the manuscript with input from all authors.

## Supplementary Figures

**Supplementary Figure 1.**
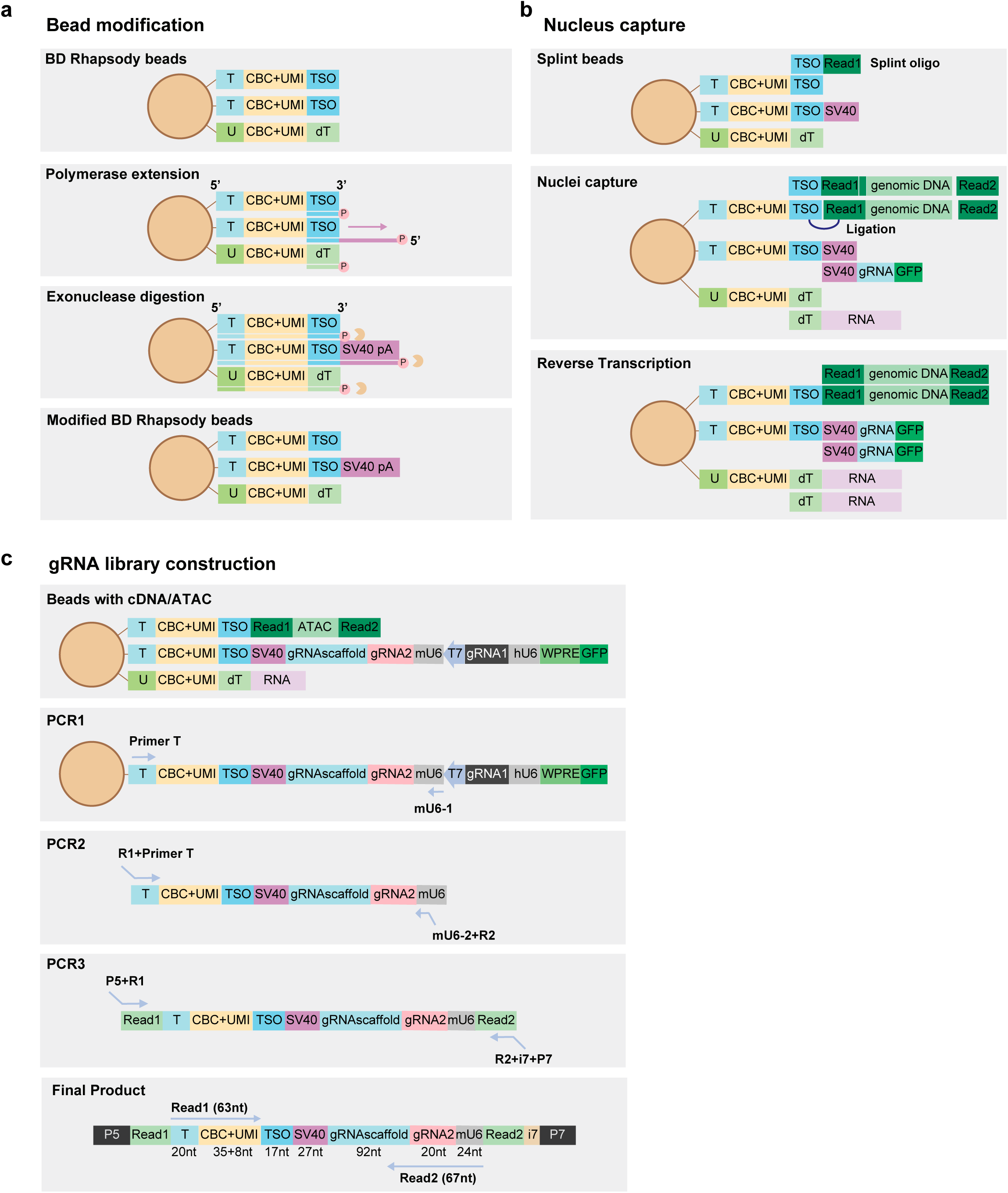
Bead modification, nuclei capture and library construction workflow for gRNA recovery in sn-multiome profiling. (a) Schematics of bead modification to introduce a gRNA-specific capture sequence onto BD Rhapsody capture beads. P in the circle indicates 5’ phosphorylation on an oligonucleotide. (b) Schematics of nucleus capture on modified beads, showing recovery of gRNA-derived transcripts together with cellular transcripts and tagmented genomic DNA fragments. (c) Nested target-enrichment (TE) PCR workflow for gRNA library construction and sequencing configuration.

**Supplementary Figure 2.**
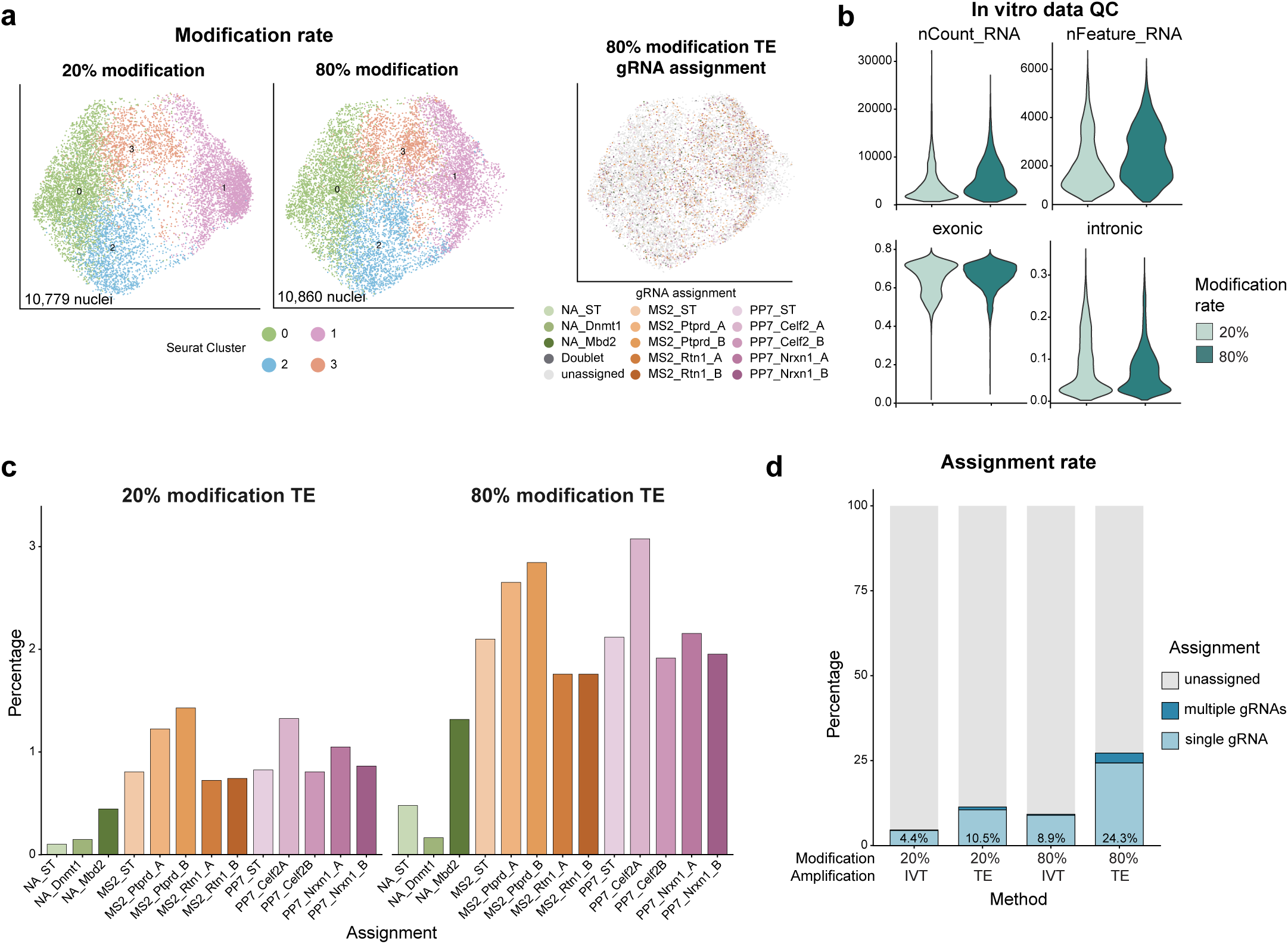
***In vitro* benchmarking of gRNA recovery across vector constructs, bead modification rates, and amplification approaches.** (a) A UMAP visualization of nuclei profiled *in vitro* at different modification rates, colored by Seurat cluster and gRNA assignment. (b) Violin plots showing RNA quality-control metrics across nuclei in the *in vitro* dataset. (c) Single-gRNA assignment rates across the vector constructs at different bead modification rates. (d) gRNA assignment rates across bead modification rates and amplification approaches.

**Supplementary Figure 3.**
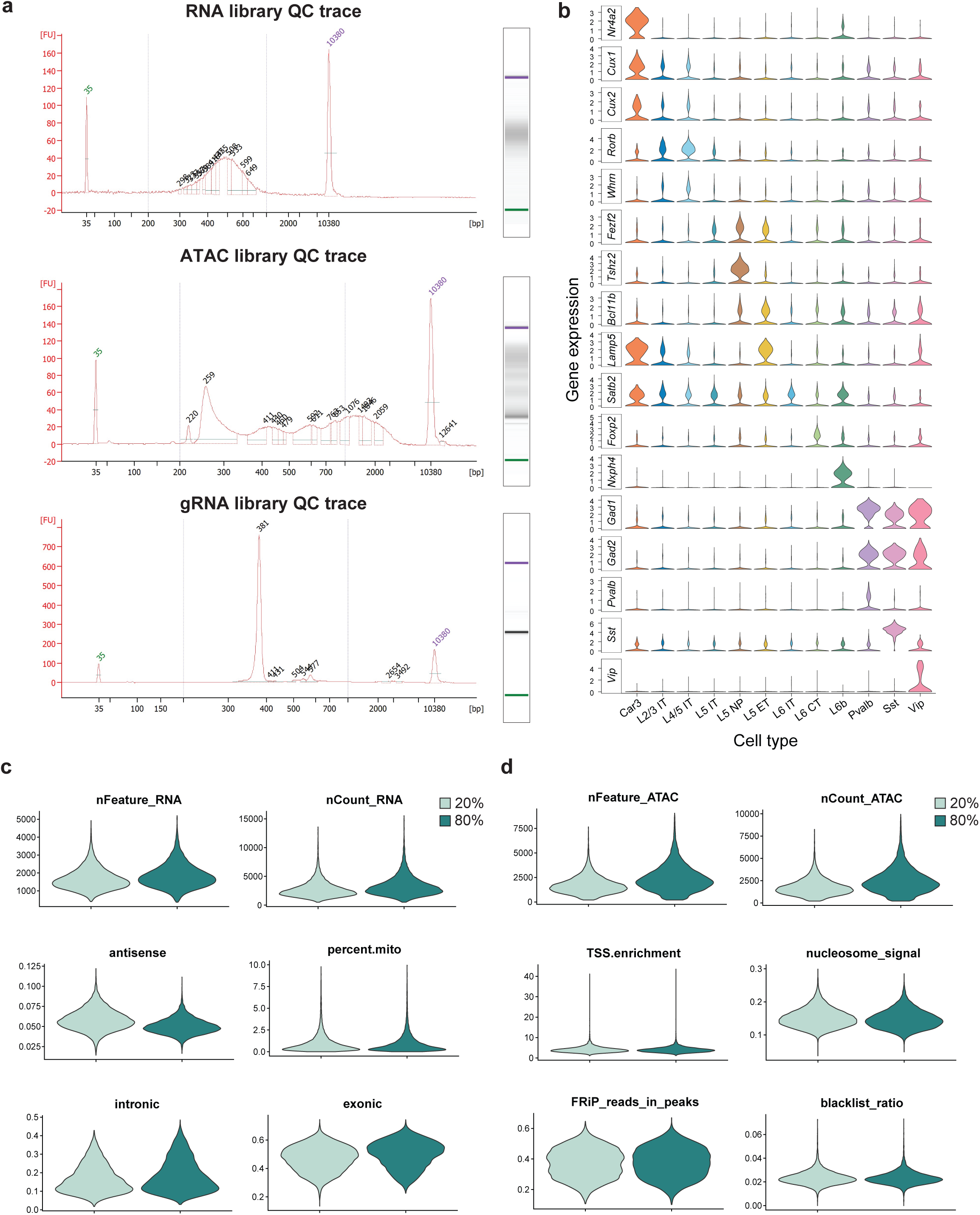
Quality assessment of the *in vivo* sn-multiome dataset. (a) Representative Bioanalyzer traces showing fragment size distributions of RNA, ATAC and gRNA libraries. (b) Violin plots showing normalized expression of representative marker genes across annotated cell types. (c-d) Violin plots showing RNA and ATAC quality-control metrics across nuclei profiled *in vivo*. nFeature_RNA: number of detected gene features per nucleus. nCount_RNA: total RNA UMI counts per nucleus. percent.mito: mitochondrial transcript fraction. nFeature_ATAC: number of accessible peaks detected per nucleus. nCount_ATAC: number of fragments per nucleus. TSS.enrichment: transcription start site enrichment score. FRiP: fraction of fragments in peaks. blacklist.ratio: fraction of ATAC molecules overlapping blacklist regions.

**Supplementary Figure 4.**
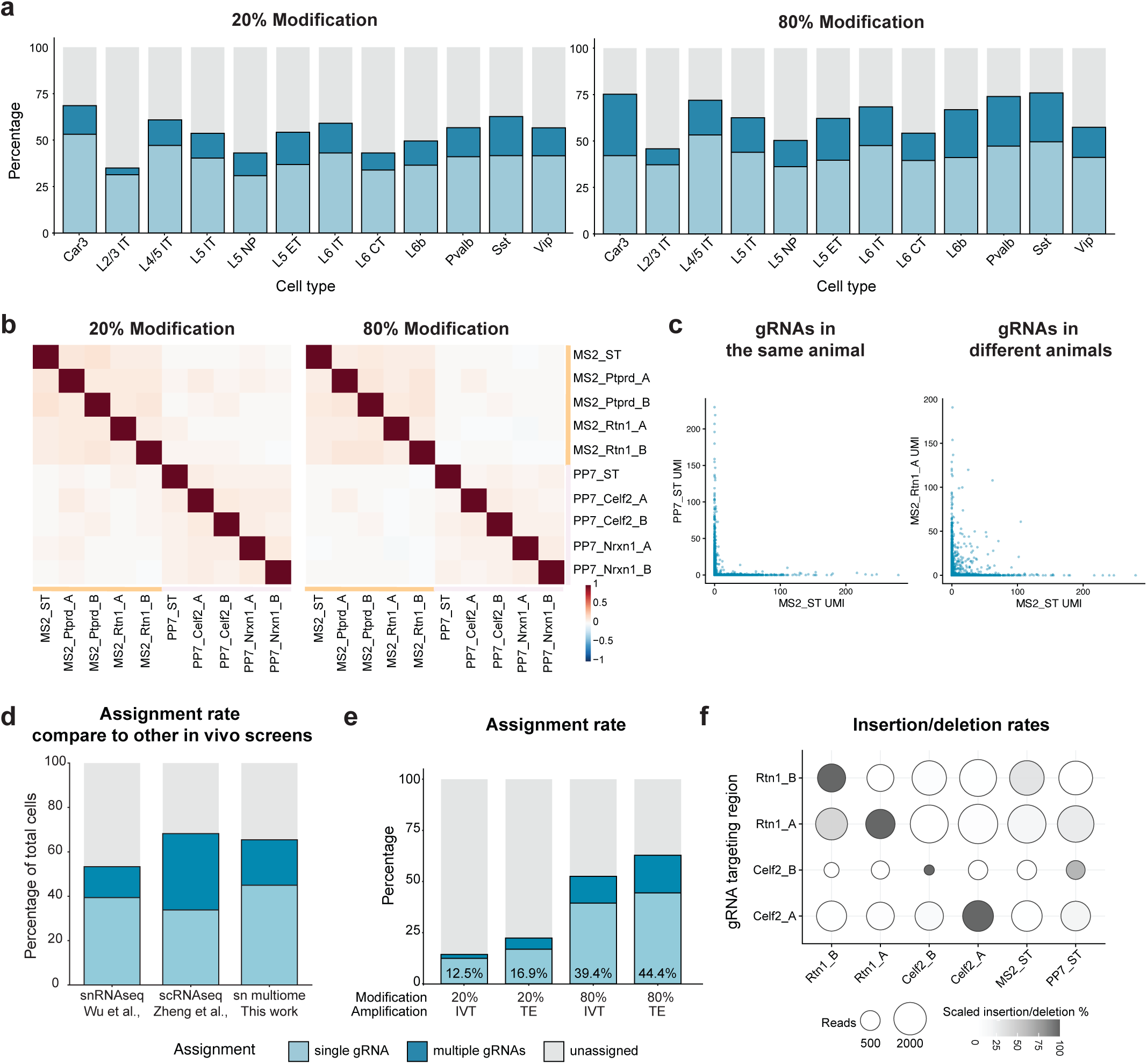
Assessment of *in vivo* gRNA assignment rate and quality. (a) gRNA assignment rates across cell types at different bead modification rates. (b) gRNA assignments across vector constructs show weak cross-gRNA correlations across bead modification rates. (c) Low cross-gRNA correlation between different gRNAs within the same animal and across animals. (d) The *in vivo* multiome Perturb-seq dataset generated in this study shows gRNA assignment rates comparable to those reported in other studies^9, 14^ with only the transcriptome. (e) gRNA assignment rates across bead modification rates and amplification approaches. (f) Dot plot showing insertion/deletion frequencies and read depth at gRNA target loci from the multiome dataset across gRNA assignments, scaled by the maximum value in each row.

**Supplementary Figure 5.**
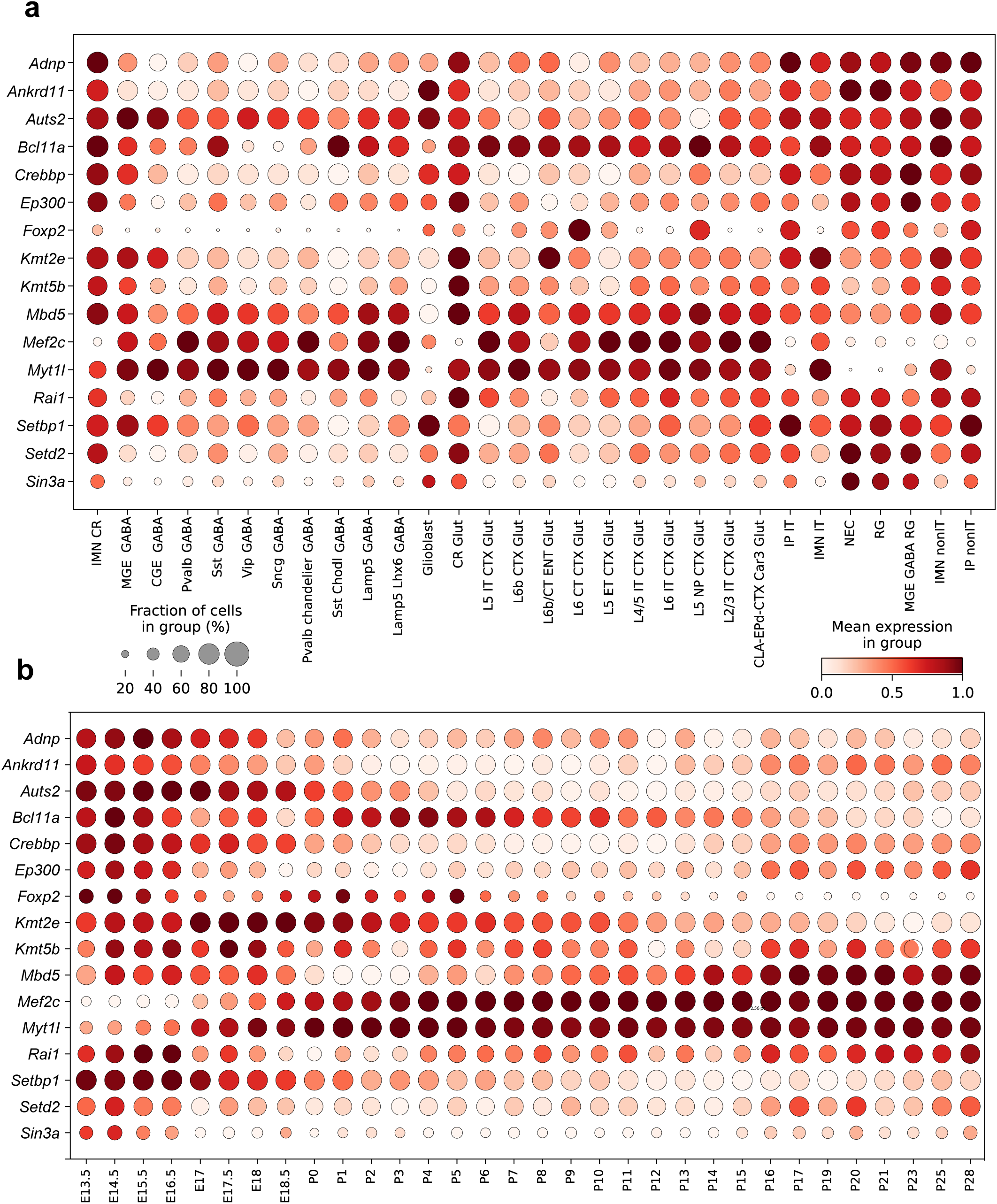
Expression of 16 NDD risk genes. (a-b) Dot plots showing expression of the 16 NDD risk genes across brain cell types (a) and developmental stages (b).

**Supplementary Figure 6.**
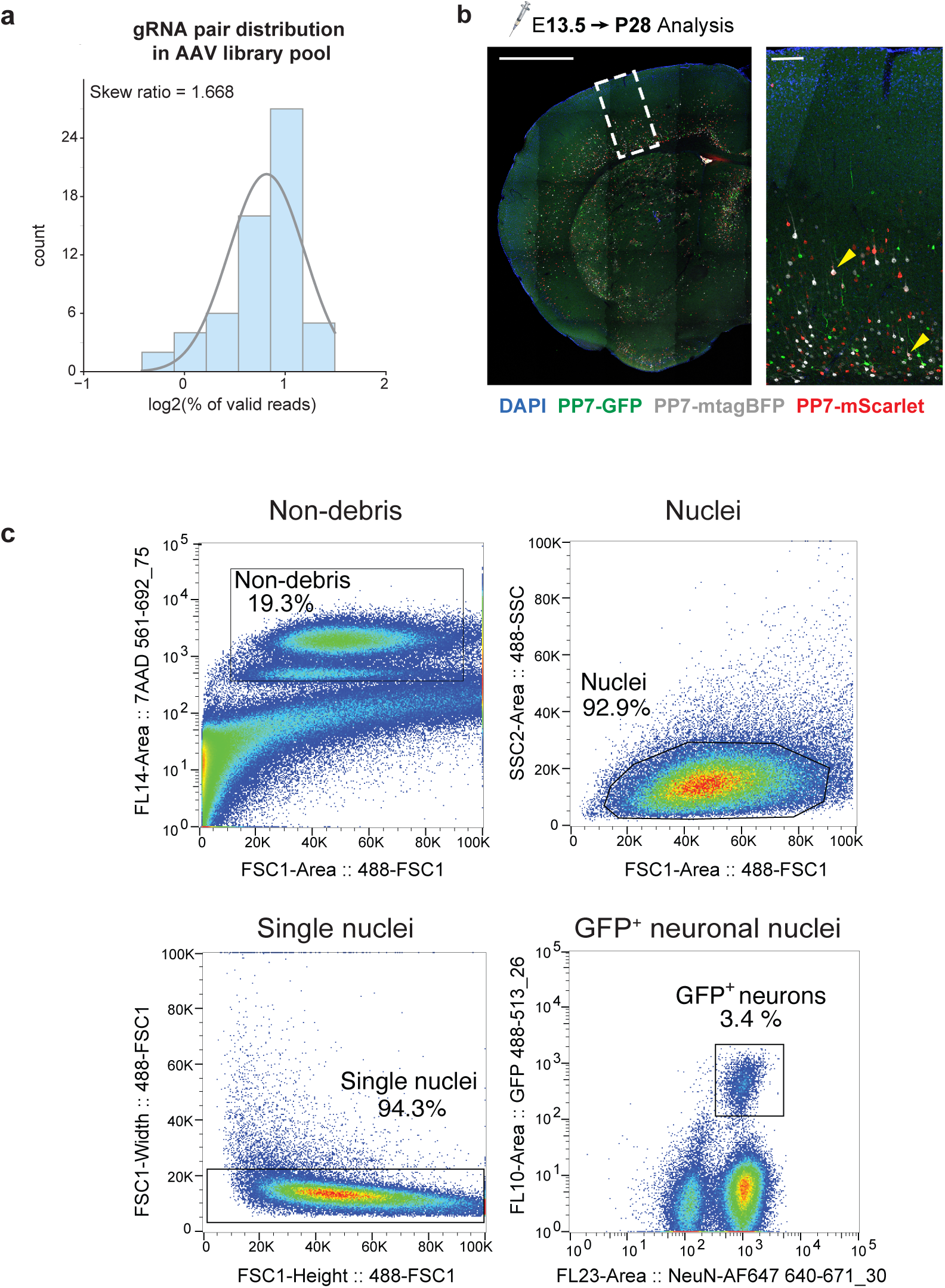
gRNA distribution in the pooled AAV library and validation of sparse AAV labeling using low MOI. (a) Distribution of gRNA pairs in the pooled AAV library determined by nanopore sequencing. Skew ratio defined as the 90th percentile divided by the 10th percentile of nonzero percentages. (b) Representative images of AAV-mediated labeling in the mouse cortex using three reporters, showing minimal co-transduction events. Yellow arrowheads indicate co-transduced cells labeled by more than one reporter (less than 1%). This viral titer at 1×10^9^ vg/embryo was optimal for maximizing the transduced cells while keeping the cells with multiple perturbations low. Scale bars 1mm and 100 μm. (c) FANS enrichment of GFP^+^ nuclei following low MOI AAV delivery.

**Supplementary Figure 7.**
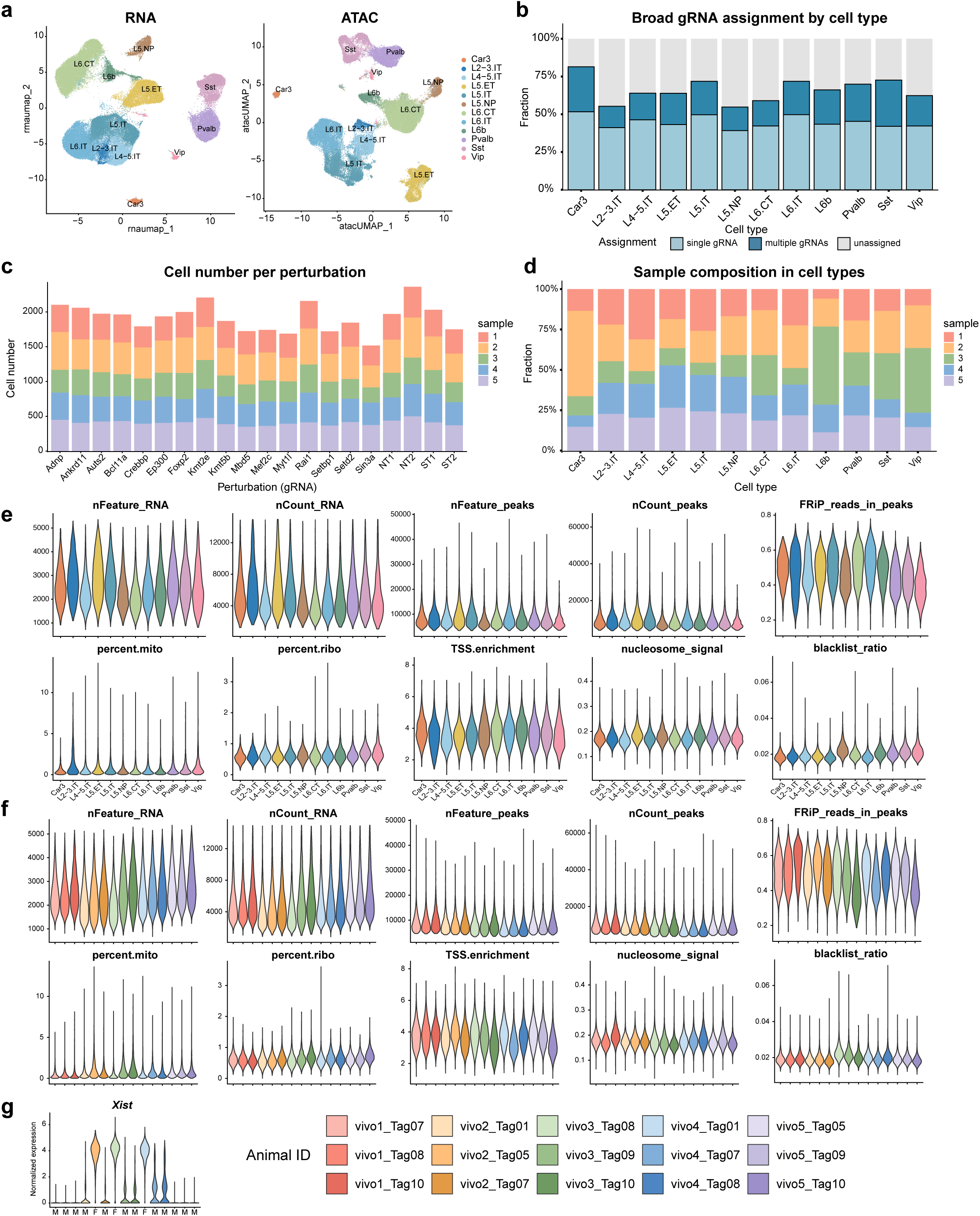
Quality assessment of RNA, ATAC, and gRNA assignments in the *in vivo* multiome Perturb-seq screen. (a) UMAP visualizations of RNA and ATAC modalities from the *in vivo* multiome dataset, colored by cell type. (b) gRNA assignment rates across annotated cell types. (c-d) Distribution of nuclei from the five samples across perturbations (c) and cell types (d). (e-f) Violin plots showing RNA and ATAC quality-control metrics across cell types (e) and animals (f), as described in Supplementary Figure 3c-d. (g) *Xist* expression in animals agreed with the sex of animals, supporting the animal identity assignment. M: male; F: female.

**Supplementary Figure 8.**
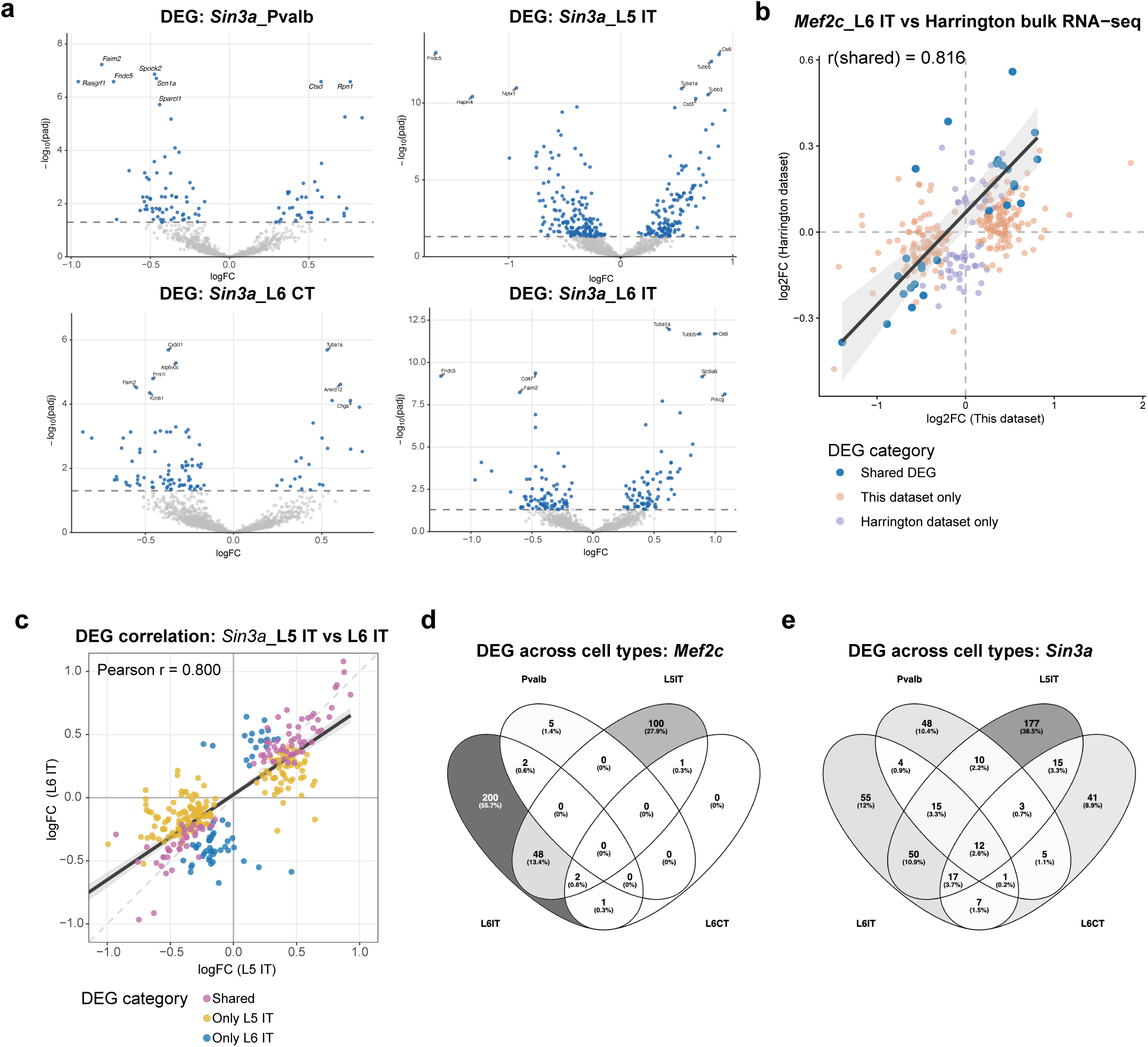
Differential gene expression and chromatin accessibility analyses across perturbed cell types. (a) Volcano plots of DE effects by *Sin3a* perturbations in Pvalb inhibitory neurons, L5 IT, L6 CT and L6 IT. (b) Scatter plot comparing DEGs by *Mef2c* perturbation in L6 IT to published RNA-seq dataset in *Mef2c* knockout mouse cortex^20^. The dark-gray line and light-gray shaded band show the linear regression fit and 95% confidence interval computed from shared genes, respectively. (c) Scatterplot comparing DEG results between L5 IT and L6 IT by *Sin3a* perturbation. The dark-gray line and light-gray shaded band show the linear regression fit and 95% confidence interval computed from all plotted genes, respectively. (d-e) Venn diagrams showing overlap of DEGs across Pvalb inhibitory neurons, L5 IT, L6 CT and L6 IT by *Mef2c* (d) and *Sin3a* (e). Color shading indicates the percentage of genes in each region relative to the total number of DEGs.

**Supplementary Figure 9.**
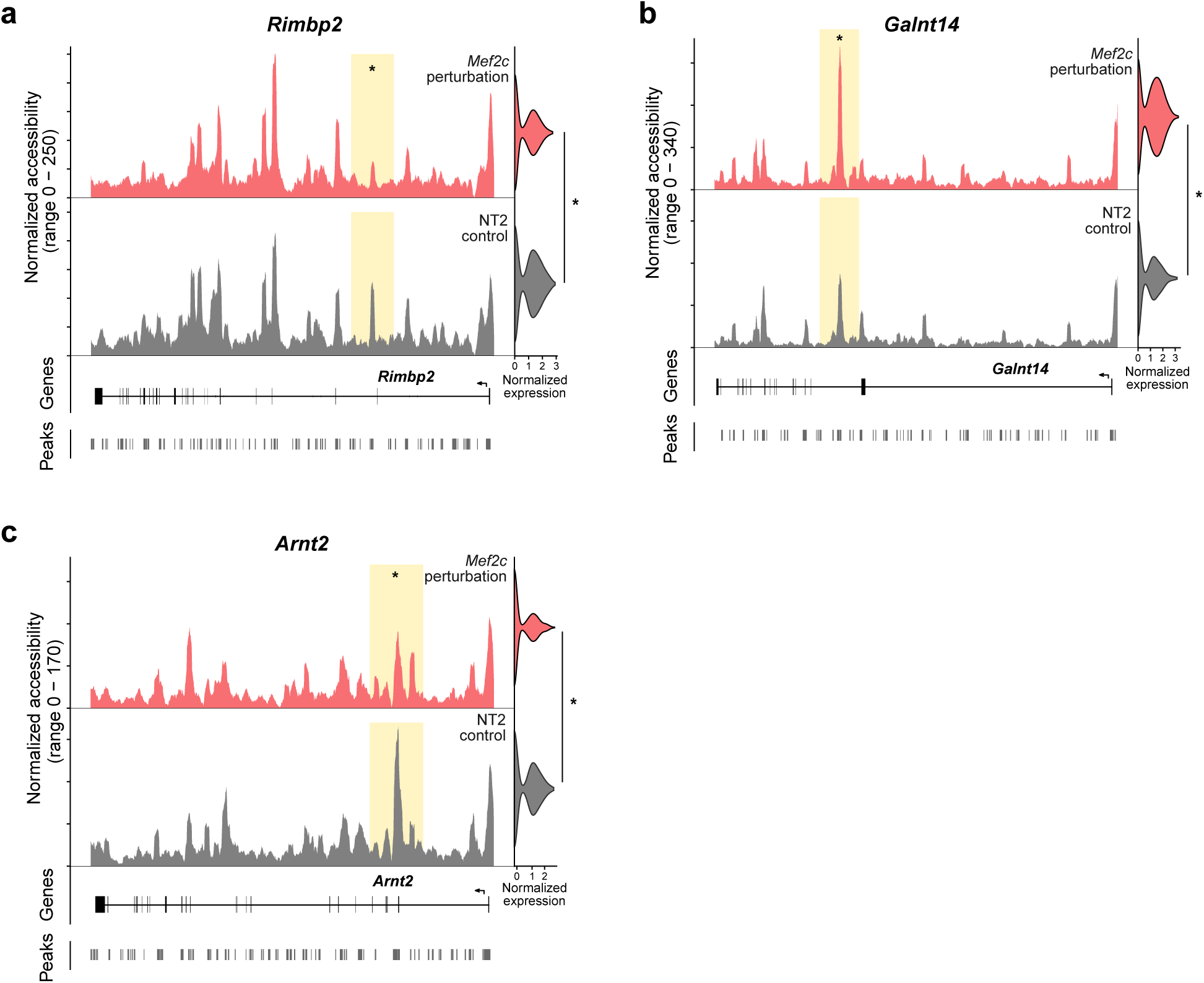
Linked chromatin accessibility and gene expression changes at specific loci in *Mef2c*-perturbed L6 IT neurons revealed by joint RNA-ATAC profiling. (a-c) Representative loci showing linked chromatin accessibility and gene expression changes in *Mef2c*-perturbed L6 IT neurons. Asterisks mark significant DAR and DEG (*P_adj_* <0.05).

## Supplementary information

Table S1. Plasmids, gRNAs, oligonucleotides and AAV vectors.

Table S2. Sn-multiome library metadata and quality metrics.

Table S3. Sn-multiome perturbation phenotypic analysis summary.

